# The power of naming: shorter and simpler species names draw more attention

**DOI:** 10.64898/2026.04.07.716944

**Authors:** Julia Mlynarek, Stephen B. Heard, Stefano Mammola

## Abstract

If you’ve ever complained about a species name that’s a mouthful—say, the soldier fly *Parastratiosphecomyia stratiosphecomyioides* or the myxobacterium *Myxococcus llanfairpwllgwyngyllgogerychwyrndrobwllllantysiliogogogochensis*—you’re in very good company. But could the readability of binomial scientific names cause more than complaints? Could it influence how much species are studied and talked about? We examined a random sample of 3,019 species names spanning 29 phyla/divisions. We tested whether name length and reading difficulty are associated with species’ representation in the scientific literature (measured via literature mentions) and their visibility to the public (measured via Wikipedia pageviews). Both species name traits showed significant negative relationships with literature mentions and Wikipedia reads. Increasing name length from 10 to 30 characters is associated with a 66% decrease in expected mentions and a 65% decrease in Wikipedia reads, while shifting from the most to the least readable name in the dataset corresponds to 53% and 76% decreases. These patterns are consistent with something familiar: the fickleness of human attention, responding to features of the world that are far from rational. While creativity in naming is a cherished part of taxonomy, a touch of orthographic restraint may ultimately benefit both science and the species themselves—especially among understudied uncharismatic taxa.

## Introduction

The names of things have power, both real (e.g. Azaryahu 1996, Wänke, Herrmann and Schaffner 2007, Graham 2011) and imagined (e.g. Guin 1968). The simple act of naming creates meaning, and influences how we understand, communicate with, and relate to the world (Wittgenstein 1922). For biologists, scientific names of species are the foundation of research and communication (Vink, Paquin and Cruickshank 2012); but the influence of species names extends beyond science into public engagement. Recent evidence suggests that characteristics and provenance of species names—such as their cultural or biological references—may shape how species are studied, cited, and recognized both inside and outside science (e.g., Heard and Mlynarek 2023, Mlynarek *et al*. 2023, Blake *et al*. 2024). For example, insect species named after host plants are more often tested for the existence of cryptic host races (Mlynarek *et al*. 2023), and species named for celebrities receive more public and media attention (Heard and Mlynarek 2023, Blake *et al*. 2024).

Scientific naming is governed by conventions formalized in the International Code of Zoological Nomenclature (ICZN 2000) for animals and the International Code of Nomenclature (ICN Turland *et al*. 2018) for algae, fungi, and plants. The Codes provide rules for forming names but grant wide latitude to taxonomists. For example, both Codes require binomial names for species in latinized form, but neither restricts the language on which a name (or name part) is based. Both codes recommend avoiding names of excessive length, but neither elevates this to a requirement; and a recommendation in the ICN that taxonomists avoid names that are difficult to pronounce has been dropped in the most recent (2025) version of the Code. Thus, while naming is regulated, the Codes leave ample room for interesting cases—as demonstrated, for example, by the astounding etymological diversity found in *Aloe* plants (Figueiredo and Smith 2010), spiders (Mammola *et al*. 2023b), phytophagous insects (Mlynarek *et al*. 2023), or parasites (Poulin, Dutra and Presswell 2022).

Several recent papers have presented best practices in naming to reduce taxonomic instability (Smith and Chiszar 2006, Ruedas, Norris and Timm 2025). Such advice highlights the potential for choices in naming to matter, and for consistent and clear naming practices to support effective communication. While there are many dimensions to how naming is done, two simple aspects of scientific names often provoke comments: length and reading difficulty. Scientific names have a reputation for being long and complex, and indeed, some are. The longest valid binomial name for any organism is *Myxococcus llanfairpwllgwyngyllgogerychwyrndrobwllllantysiliogogogochensis*, for a myxobacterium named after the Welsh village with Europe’s longest place name (Chambers *et al*. 2020). Among animals, the record belongs to the soldier fly *Parastratiosphecomyia stratiosphecomyioides* (Brunetti 1923). While these names are valid under their respective Codes, they are arguably unfortunate and illustrate the absence of enforceable constraints on length. In rare cases, excessive length and reading difficulty has led to suppression of names for practical reasons; for instance, a set of amphipod names including *Gammaracanthuskytodermogammarus loricatobaicalensis* were suppressed under Suspension of the Rules by the ICZN because the alternative was “greater confusion than uniformity” (ICZN and Hemming 1929). Such incidents highlight tension between taxonomic freedom and usability.

We investigated whether name length and reading difficulty are associated with species’ representation in the scientific literature and with their visibility to the public (the latter estimated via Wikipedia page views). Such an association could arise because longer, difficult-to-read names increase cognitive load (New *et al*. 2006); for this reason we hypothesized that shorter, more readable names would garner, on average, more scientific and societal attention. On the other hand, it is conceivable that *longer* names might garner more attention because they are unusual thus capture attention. By testing these hypotheses, we aim to contribute to a growing body of work that examines how taxonomic decisions may influence the broader impact and accessibility of biodiversity knowledge.

## Material & Methods

We used a dataset of 3,019 species names (across 29 phyla/divisions) (Mammola *et al*. 2023a) to test the effect of species name length and overall readability on mentions of the species in the scientific literature and on views of species Wikipedia pages. While the dataset was developed to study the influence of species-level traits on scientific and societal attention, we repurposed it to test our working hypothesis. This dataset was compiled through random stratified sampling of the eukaryotic multicellular Tree of Life, encompassing Animalia, Fungi (restricted to Agaricomycetes), and Plantae (excluding unicellular algae). The species in this dataset were named between 1758 and 2019. The dataset includes the number of articles indexed in the Web of Science that refer to each species, which provides a quantitative estimate of research effort devoted to individual species and hence a proxy measure of scientific interest (Wilson *et al*. 2007). Additionally, for each species, the dataset reports the total number of Wikipedia pageviews across all languages in which the species is represented, serving as a proxy for general societal interest in that species (Mittermeier *et al*. 2021). Further details about the dataset and data collection methods can be found in Mammola *et al*. (2023a).

For each species, we calculated the length of the species binomial name (i.e., the sum of genus and species) and an index of name readability (Figure S1). We estimated the latter with the following formula:

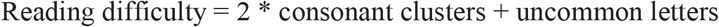

The index integrates two features of the name that can influence visual and phonetic processing effort: i) the number of consonant clusters of three or more letters (e.g., *“str”, “phl”*), which tend to reduce pronounceability and readability; and ii) the number of uncommon letters (*j, k, x, y*, and *z*), given that these letters are relatively rare in Latinized nomenclature and may reduce familiarity. The consonant cluster component is weighted to reflect its higher contribution to phonetic complexity. Name length and reading difficulty were positively correlated (Pearson’s *r* = 0.556, *t*_(3017)_ = 36.7, *p* < 2.2 × 10^−16^), as longer names tend to contain, on average, more clusters and uncommon letters than shorter names. Consequently, we regressed reading difficulty on length and extracted the model residuals, where positive values indicate species names that are harder to read, and negative values indicate names that are easier to read, than expected for their length. We recognize the impossibility of measuring reading difficulty in a culturally universal way and return to this point in the Discussion.

We tested for effects of length and reading difficulty on attention using regression analyses (Zuur and Ieno 2016). We ran all analyses in R version 4.4.1 (R Core Team 2024), using the package *glmmTMB* version 1.1.10 for modelling (Brooks *et al*. 2017), *performance* version 0.12.4 for model validation (Lüdecke *et al*. 2021), and *ggplot2* version 3.5.1 for visualizations (Wickham 2016). Specifically, we fitted two generalized linear mixed models to test the effects of name length and reading difficulty on (i) literature mentions and (ii) Wikipedia pageviews. Because both response variables are counts, we initially assumed a Poisson error structure. However, as both Poisson models showed overdispersion, we refitted them using a negative binomial distribution—a generalization of the Poisson model that relaxes the assumption that the variance equals the mean. Each model included the year of species description as an additional covariate to account for the fact that older species have had more time to accumulate literature mentions and Wikipedia pageviews. To account for taxonomic non-independence, we included a nested random intercept structure (1 | Phylum / Class / Order), assuming that closely related species may have more similar names and levels of human interest than expected by chance.

We acknowledge that literature mentions and online attention patterns are influenced by multiple factors beyond those accounted for in our regression analysis. These include not only scientific considerations and species-level characteristics (such as the ones analyzed in Mammola *et al*. (2023a), but also publication-level factors including author visibility, journal prestige, open access, and related elements (Tahamtan, Safipour Afshar and Ahamdzadeh 2016, Tahamtan and Bornmann 2019, Mammola *et al*. 2022). However, we chose to keep the analysis focused on the species-name characteristics of interest (length and readability); the total variance explained by both our models (R^2^ < 0.3; see Results) likely reflects the omission of these additional factors. We note that in an alternative analysis including all factors considered by Mammola *et al*. (2023a), name length and readability remain significant.

## Results

Unsurprisingly, both literature mentions and Wikipedia pageviews are extremely variable across species (mentions, range 0 – 7280, median 0, SD 217; pageviews, range 0 – 5 × 10^7^, median 269, SD 1.4 × 10^6^). Both literature mentions (p<0.0001; Table S1) and Wikipedia reads (p<0.0001; Table S2) increase significantly with time since publication, justifying our decision to include publication year as a predictor in our models. The shortest names in our dataset were *Crex crex, Ninox ios*, and *Puda puda* (all 8 letters); the longest were *Callorhynchicola multitesticulatus, Demicryptochironomus cinereithorax*, and *Saemundssonia scolopacisphaeopodis* (all 33 letters). After correction for length, the most readable names in our dataset were *Alosa alosa, Isoetes sabatina*, and (perhaps counterintuitively) *Unionicola inusitata*. The least readable were *Amphiastrella kirkpatricki, Polyacanthorhynchus kenyensis*, and *Pseudoscaphirhynchus fedtschenkoi*, and we apologize for this sentence.

Controlling for year of description, species with longer or less readable names tended to receive less scientific and societal attention (Figure 1; see Table S1–S2 for exact model estimates). Specifically, both name length and reading difficulty showed significant negative relationships with literature mentions (Figure 1A–B) and Wikipedia pageviews (Figure 1C–D). ***[See figures next page*.]** While the data are unsurprisingly noisy (our models account for 29% and 13% of variance in literature mentions and Wikipedia pageviews, respectively), the effects are not small. For instance, increasing name length from 10 to 30 characters is associated with a 66% decrease in expected mentions and a 65% decrease in Wikipedia pageviews, while reducing the reading difficulty index from the most readable name to the least readable species name in the dataset corresponds to 53% and 76% decreases.

**Figure 1:**
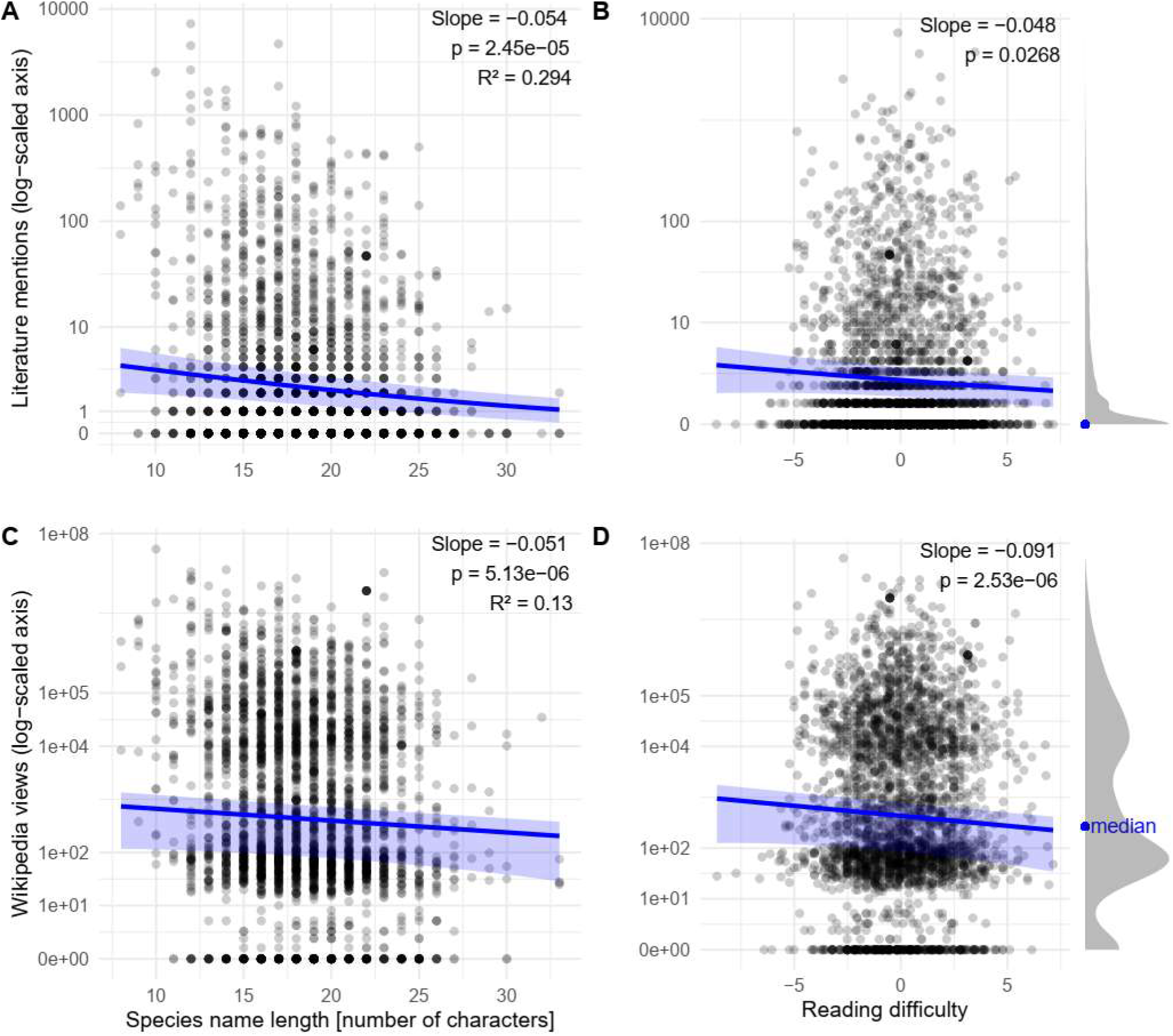
Relationships between species name length, public attention, and reading difficulty across taxa. (A) Relationship between species name length (number of characters) and the number of scientific literature mentions (log-scaled). (B) Relationship between species name length and Wikipedia page views (log-scaled). (C) Association between species name length and the reading difficulty of the corresponding Wikipedia entry. (D) Median distribution of reading difficulty scores. Slopes, p-values, and R^2^ values for each linear model are shown. Density curves to the right of each pair of plots show data distribution for literature mentions and Wikipedia pageviews, with blue dots marking the median values.

## Discussion

We find that species with longer and less readable names receive less attention from scientists and from the public. We are not, of course, suggesting that anyone *deliberately* ignores species with longer names. Instead, we suspect that (assuming the effect is causal) unconscious reactions to long, complex names are responsible. This simple observation underscores that human attention is fickle and can respond to features of the world in ways that are far from rational. After all, there is to our knowledge no evidence that species with longer or less readable names are less important ecologically, less in need of conservation protection, or less fascinating in their morphology and behaviour.

There is other recent evidence that the decisions taxonomists make in coining new species names can have surprising implications for later study of those species. Mlynarek *et al*. (2023) focused on the etymology of names, showing that among plant-feeding insects, those named after their host plants are more than twice as likely to later be the subject of studies testing for the existence of cryptic host-specialist races. Here our question was more general: do simple properties of species names affect the frequency with which they are studied *at all?* (They do.) There are, of course, many other considerations that influence scientists’ choice of study organisms (e.g. Westoby 2002, Dietrich *et al*. 2020, Adamo *et al*. 2021). Some of these are entirely rational: species with economic importance or under conservation threat may receive considerable scientific attention (Mammola *et al*. 2023a). Others are less so: species widely considered charismatic (vertebrates) receive outsized attention (Troudet *et al*. 2017), species in inaccessible habitats (Ficetola, Canedoli and Stoch 2019), and very small species may be understudied for practical reasons (Vásquez-Restrepo 2026). It would be difficult, though, to conceive of a reason for attention (or lack of it) as unrelated to scientific needs as the length and readability of the species’ scientific name. Scientists are, it appears, entirely human in their idiosyncrasy.

It is perhaps less surprising that public attention to a species might be reduced by the barrier posed by a long, difficult name. Every biologist has heard non-scientists (and biology students, for that matter) objecting to scientific names as generally impossible to remember or pronounce. These objections are not entirely ill-founded: consider, for example, the long and convoluted bacterium name *Thermodesulfobacterium hydrogeniphilum*, or the phonetic ordeal in the spider *Chilobrachys jonitriantisvansickleae*. Of course, for species that have a common name, members of the public are likely to use that instead, and this may be partly why we found weaker effects for Wikipedia pageviews than for mentions in the scientific literature. This hypothesis would be straightforward to test with a large enough dataset of Wikipedia articles that do and don’t include common names for the species they cover. However, across the tree of life most species do not have common names (or their common names are just anglicizations of their scientific names). For such species, any public attention they receive will necessarily come via their scientific names.

Length and reading difficulty aren’t the only features of names that may influence public attention, of course. Similarly, Blake *et al*. (2024) and Heard and Mlynarek (2023) found that species named after celebrities get, on average, more online and mass-media attention— making this a naming strategy that could be particularly significant for taxonomic groups that are generally less popular than others (e.g., invertebrates, fungi).

Human reactions to features of names may influence more than just attention. Two studies have asked whether species names influence public interest in conservation efforts for those species. Karaffa, Draheim and Parsons (2012) found that names with positive emotional resonance (e.g., “Patriot Eagle”) attracted more conservation interest from university students than those with negative emotional resonance (e.g., “Razor Eagle”). In this case, the names were uninformative about the species but still had semantic implications. Díaz-Restrepo *et al*. (2022), in contrast, examined strictly orthographic aspects of names, comparing (invented) common names that were short and pronounceable and that began with plosive consonants (B, C, D, K, G, P or T) versus those that were not. Each of these features is associated with effective product brand names (e.g., Bergh *et al*. 1984, Bao, Shao and Rivers 2008). They found no effect of name on conservation interest from adults in the United Kingdom, although they did not attempt to isolate effects of length from those of initial letter choice. The large literature on construction of product brand names—and the strong but complex human preferences it reveals—suggests that further work along these lines would be rewarding.

While name length is easily measured, we acknowledge that our measure of readability focuses on a feature (consonant clusters) that might be considered more difficult for speakers of some languages (e.g., Romance, Semitic, Sinitic, and Austronesian languages) than for others (e.g., Slavic and Kartvelic languages). Similarly, the letters we have scored as “uncommon” in Latinized nomenclature are quite common in some languages. Given extreme differences in phonological makeup among the world’s 7,000 or so extant languages, it is likely not possible to design a universal measure of readability. Still, the significant role our readability score plays in predicting both literature mentions and Wikipedia pageviews suggests that it’s capturing something informative. We speculate that this may be because aversion to consonant clusters and to the letters we designate “uncommon” happens to be shared by several language families that are overrepresented among practitioners of so-called Western science (Romance, Sinitic, and Japonic, and to a lesser degree English). A first approach to the non-universality of “readability” might be to code literature mentions by the presumed language background of authors, or to break down Wikipedia pageviews by article language (and thus by presumed language background of readers). Meanwhile, it would be interesting to consider the effects of other linguistic features that might make names more or less readable (or memorable), such as strings of visually similar letters or the occurrence of rare letter combinations rather than rare single letters.

Going beyond readability, our analysis could be extended to consider other features of name morphology. Since etymology had a strong effect on a particular aspect of scientific attention in Mlynarek *et al*.’s (2023) study, it would also be interesting to ask whether species varying in name etymology (those named for morphology, habitat, geography, or eponymously; (Heard and Mlynarek 2023)) receive differing scientific attention more generally. In particular, we wonder whether species named using jokes or cultural references (e.g., the tiny frogs *Mini mum, Mini scule, and Mini ature*; (Scherz *et al*. 2019); see also https://www.curioustaxonomy.net) might receive more or less attention as a consequence. While humour and cultural references in science are sometimes derided, their inclusion in paper titles increases citation rates (at least in ecology and evolution (Heard, Cull and White 2023)). A similar effect for species names, if it exists, would be an intriguing parallel.

Our findings suggest that the act of naming species carries weight. Taxonomists might, thus, consider the likely effects of their naming choices on the users of those names—in this case, scientists who research the species or members of the public curious about them. For example, a taxonomist might choose species names strategically to increase their visibility (particularly useful for understudied “dark taxa”). However, naming decisions might also have the opposite effect, inadvertently hiding a species from attention it might otherwise receive. Our results also imply that suggestions in the International Codes of Botanical and Zoological Nomenclature to avoid very long names (Botanical, Recommendation 23A.3(b), (Turland *et al*. 2018); Zoological, Recommendation 25C, (ICZN 2000)) are well founded. Although recommendations about pronounceability have recently been deleted from the Botanical Code and are only implied in the Zoological code, our results suggest that taxonomists might nevertheless benefit from considering pronounceability, which is closely connected to readability.

Taking a different perspective, the ways taxonomists have historically named species have sometimes perpetuated systemic and societal biases (e.g., gender disparities or echoes of colonialism; (Guedes *et al*. 2023, Chala *et al*. 2024, Pétillon *et al*. 2025)). This is relevant here because sometimes, a taxonomist might choose a name that references, or pays homage to, a region, culture, or language appropriate to the species being named. The result might be a name that, because it draws on Japanese, Georgian, or Xhosa linguistic and orthographic patterns, is more difficult for many or even most global scientists. A consequent reduction in scientific and public attention to the species might be considered an acceptable trade-off. However, taxonomists will be better equipped to consider such trade-offs if they are awareness of the kind of attention effect we document here.

In the end, no-one should be surprised to find that naming choices matter. We do not wish to suggest that taxonomists refrain from creativity, or from cultural sensitivity, in naming; however, tempering that creativity with some orthographic caution may ultimately benefit both science and the species being named.

## Data and code availability

Data supporting this study are available in Figshare (https://doi.org/10.6084/m9.figshare.22731440) and fully described in Mammola et al. (2023a). Annotated code to reproduce analyses is available on GitHub (https://github.com/StefanoMammola/Analysis_name_length)

## Notes

https://doi.org/10.6084/m9.figshare.22731440

https://github.com/StefanoMammola/Analysis_name_length

### Competing Interest Statement

The authors have declared no competing interest.

