## Supplemental tables and figures for "The power of naming: shorter and simpler species names draw more attention"

**Supporting information**

**Table S1. Estimated parameters for the regression model with citations as the dependent variable.**

| **Variable** | **Estimate** | **Standard Error** | **z** | **p** |
|---|---|---|---|---|
| Intercept | 33.87 | 1.46 | 23.08 | - |
| Year of publication | –0.01 | 0.0007 | –22.38 | $<2e^{-16}$ |
| Name length | –0.05 | 0.01 | –4.22 | $2.45e^{-5}$ |
| Reading difficulty | –0.04 | 0.02 | –2.21 | 0.026 |

**Table S2. Estimated parameters for the regression model with Wikipedia as the dependent variable.**

| **Variable** | **Estimate** | **Standard Error** | **z** | **p** |
|---|---|---|---|---|
| Intercept | 33.45 | 1.31 | 25.48 | - |
| Year of publication | –0.01 | 0.0006 | –21.58 | $<2e^{-16}$ |
| Name length | –0.05 | 0.01 | –4.55 | $5.13e^{-6}$ |
| Reading difficulty | –0.09 | 0.01 | –4.70 | $2.53e^{-6}$ |

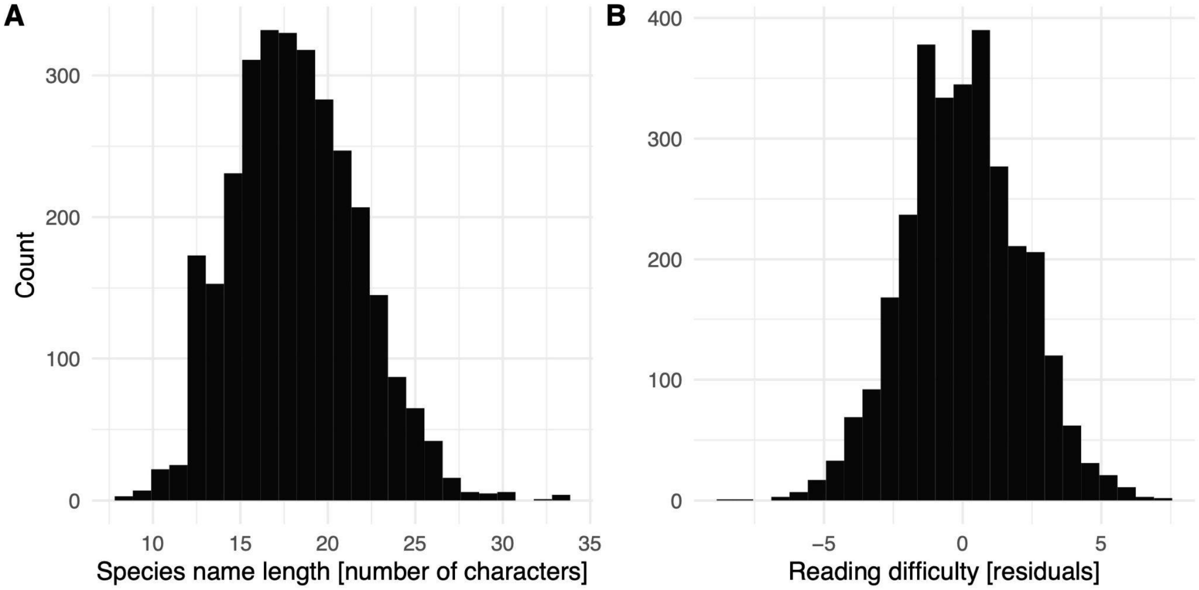


**Figure S1. Distribution of species name length and readability.** Histograms showing (**A**) the distribution of species name lengths (in number of characters) and (**B**) the distribution of name readability scores across all species in the dataset (expressed as residuals of a regression model of readability scores against length of the name).
